# Sex-specific gut microbiota modulation of aversive conditioning and basolateral amygdala dendritic spine density

**DOI:** 10.1101/2020.07.21.213116

**Authors:** Caroline Grace Geary, Victoria Christina Wilk, Katherine Louise Barton, Parvaneh Ottavia Jefferson, Tea Binder, Vasvi Bhutani, Claire Baker, Achal James Fernando-Peiris, Alexa Lee Mousley, Stefano Freitas Andrade Rozental, Hannah Mae Thompson, Justin Charles Touchon, David Justin Esteban, Hadley Creighton Bergstrom

## Abstract

Gut microbiota influence numerous aspects of host biology, including brain structure and function. Growing evidence implicates gut microbiota in aversive conditioning and anxiety-related behaviors, but research has focused almost exclusively on males. To investigate sex-specific effects of gut dysbiosis on aversive learning and memory, adult female and male C57BL/6N mice were orally administered a moderate dose of non-absorbable antimicrobial medications (ATMs; neomycin, bacitracin, pimaricin) or a control over 10 days. Changes in gut microbiome composition were analyzed by 16S rRNA sequencing. Open field behavior, cued aversive learning, context recall, and cued recall were assessed. Following behavioral testing, the morphology of basolateral amygdala (BLA) principal neuron dendrites and spines was characterized. Results revealed that ATMs induced distinct but overlapping patterns of gut dysbiosis across sex, with stronger effects in females. There were also sex-specific effects on behavior and neuroanatomy. Treated males but not females exhibited altered locomotor and anxiety-like behavior in the novel open field test. Treated females but not males showed impairments in aversive memory acquisition and cued recall. Context recall remained intact in both sexes, as did dendritic structure of BLA principal neurons. However, ATMs exerted sex-specific effects on spine density. A second experiment was conducted to isolate gut perturbation to cued recall. Results revealed no effect of ATMs on recall of a previously consolidated fear memory, suggesting that gut dysbiosis preferentially impacts aversive learning. These data shed new light on how gut microbiota interact with sex to influence aversive conditioning, anxiety-like behavior, and BLA dendritic spine architecture.

**Significance:** Gut microbiota can influence brain function and behavior, including trauma and anxiety-related disorders. Although these disorders disproportionately affect women, preclinical research has focused almost exclusively on male rodent models. We investigated the impact of antimicrobial administration on gut microbiome structure, aversive conditioning, open field behavior, and basolateral amygdala principal neuron morphology in female and male mice. Results showed that treatment exerted wide-ranging effects, many of which were sex-specific. Our findings underscore the importance of studying sex differences and support a role for microbial modulation of aversive learning, anxiety-like behavior, and amygdala spine patterning.

## 1 INTRODUCTION

The gastrointestinal tract harbors a diverse microbial ecosystem which plays a fundamental role in human health. Although the ability of pathogens to modulate host physiology and behavior is well established (Lyte, 2013), important influences of resident microbiota did not enter the spotlight of neuroscience until Sudo et al. (2004) found that microbiota modulate neural networks involved in stress responsivity. Since then, a surge of evidence has amassed linking resident microbiota with host behavior and brain function, including processes such as synaptogenesis, myelination, tryptophan metabolism, microglial activation, and regulation of neurotransmitters and neurotrophic factors (Abdel-Haq, Schlachetzki, Glass, & Mazmanian, 2019; Bercik et al., 2011; Chu et al., 2019; Cryan & O’mahony, 2011; Desbonnet et al., 2015; Heijtz et al., 2011). Gut microbiota components and metabolites can act both locally and systemically through neural, immune, and endocrine pathways of communication between the gut and the brain—now termed the microbiota-gut-brain (MGB) axis—which comprise a bidirectional communication network (Cryan & Dinan, 2012; El Aidy, Stilling, Dinan, & Cryan, 2016; Forsythe, Kunze, & Bienenstock, 2016; Martin, Osadchiy, Kalani, & Mayer, 2018).

Gut dysbiosis (broadly defined as disruption of the homeostatic balance between host and associated intestinal microbiota, and characterized by a substantial shift in bacterial composition or metabolic activities) is associated with many chronic illnesses, particularly functional gastrointestinal disorders like irritable bowel syndrome (IBS; DeGruttola, Low, Mizoguchi, & Mizoguchi, 2016; Kawoos et al., 2017; van de Guchte, Blottière, & Doré, 2018). Prevalence of anxious and depressive symptoms in patients with functional bowel disorders is high, relative to both the general population as well as patients with other chronic illnesses (Jones, Latinovic, Charlton, & Gulliford, 2006; MacQueen, Surette, & Moayyedi, 2017). Increasing evidence suggests that the MGB axis may be a therapeutic target for a wide range of psychiatric conditions, including trauma and anxiety-related disorders (Cowan et al., 2018; Jaggar, Rea, Spicak, Dinan, & Cryan, 2019; Malan-Muller et al., 2018; MacQueen, Surette, & Moayyedi, 2017).

A leading behavioral model for associative “fear” learning and memory is aversive conditioning (Bergstrom, 2016). In classical aversive conditioning, a sensory stimulus, such as an auditory tone, is temporally paired with a naturally aversive stimulus, such as a mild foot shock (the unconditioned stimulus, US). Following learning, the tone by itself (now a conditioned stimulus, CS) is capable of triggering a conditioned defensive response, such as freezing. Studies generally assess subjects’ freezing response to a discrete CS (“cued recall”) and, separately, to the testing chamber in which learning occurred (“context recall”). Studies also commonly assess the ability to learn when a CS no longer predicts an associated US (“extinction”).

Various probiotics (microbiota beneficial to health) have been found to enhance aspects of aversive conditioning in male rodents, including learning, cued and context recall, and extinction (Bravo et al., 2011; Fox et al., 2017; Savignac, Tramullas, Kiely, Dinan, & Cryan, 2015). In contrast, germ-free (GF) male mice exhibit impaired cued aversive memory recall and extinction (Chu et al., 2019; Hoban et al., 2018). Treatment with antimicrobial drugs (ATMs) is another well-established method of disrupting gut microbiota homeostasis, and has important clinical relevance given the widespread global use of antibiotics (Bercik et al., 2011; Jaggar et al., 2019; Rousham, Unicomb, & Islam, 2018). Both ATM-treated and gnotobiotic (colonized by a small consortium of known bacteria) male mice have been found to exhibit impaired aversive extinction (Chu et al., 2019), suggesting that, even in developed adult brains, normal aversive conditioning circuitry depends on continuous input from a rich and diverse gut microbiome.

The amygdala is a central structure in emotional processing, motivation, and associative learning (LeDoux, 2000). Considerable research implicates the amygdala in MGB signaling: altered or absent microbial communities have been linked to changes in amygdala transcriptome (Hoban et al., 2016; 2017; 2018), neurochemistry (Bercik et al., 2011; Bravo et al., 2011), and structure (Luczynski et al., 2016; see Cowan et al., 2018 for a review). Morphological changes in amygdala principal neuron dendrites and spines—key indicators of experience-induced plasticity (Chklovskii, Mel, & Svoboda, 2004)—have also been reported in GF male mice (Luczynski et al., 2016). Together, these data support a model in which gut microbiota modulate aversive conditioning and amygdala neuroplasticity.

Although a role for the MGB axis in aversive conditioning and amygdala plasticity has been established, preclinical research has centered almost exclusively on male rodents. Considering that women are more likely to be diagnosed with a trauma or anxiety-related disorder than men (Altemus, Sarvaiya, & Epperson, 2014), the study of sex differences in aversive conditioning and anxiety-related behaviors may have important implications for psychiatric care. Sex differences in aversive conditioning (Asok, Kandel, & Rayman, 2019; Blume et al., 2017) and gut microbiome structure (Jašarević, Morrison, & Bale, 2016; Markle et al., 2013) have also been reported, albeit inconsistently (Ding & Schloss, 2014; Org et al., 2016). Furthermore, functional bowel disorders like IBS, the most common form, disproportionately affect women (Kibune-Nagasako, García-Montes, Silva-Lorena, & Aparecida-Mesquita, 2016; Lovell & Ford, 2012).

To address a research gap in the characterization of sex differences related to microbial modulation of aversive conditioning, we induced gut dysbiosis in female and male C57BL/6N mice using a moderate impact ATM cocktail administered in drinking water for 10 days. We assessed cued and contextual aversive learning and memory, as well as locomotor activity and anxiety-like behavior in the novel open field test. 16S rRNA sequencing was used to characterize shifts in gut bacterial community structure. Following behavioral testing, we analyzed changes in architecture of basolateral amygdala (BLA) principal neuron dendrites and dendritic spine density. Because treatment impaired both aversive learning and cued recall in female mice, we conducted a second, smaller-scale experiment to isolate effects of gut dysbiosis to cued aversive memory retrieval. In this experiment, ATMs were administered after consolidation of the cued aversive memory, and extinction learning and estrous stage were also examined.

## 2 METHODS AND MATERIALS

### 2.1 Animals

Female (n = 46; weight = 29.0 g ± 0.8 (mean ± SEM, here and throughout)) and male (n = 48; weight = 29.2 g ± 0.7) C57BL/6N (B6) mice were 10–16 weeks-old (mean = 13 weeks) at the beginning of the first experiment. Female (n = 24; weight = 26.5 g ± 0.8) and male (n = 24; weight = 33.8 g ± 0.8) B6 mice were 10–16 weeks-old (mean = 14 weeks) at the beginning of the second experiment. Mice were originally derived from Charles River Laboratories (Wilmington, MA) and bred in-house at the Vassar College vivarium (Poughkeepsie, NY) over multiple generations. Each year mice were supplemented into the breeding colony to maintain genetic heterogeneity. Mice were kept on a 12-h light/dark cycle (lights on at 0600) in standard polycarbonate cages (28 cm × 17 cm × 12 cm) in a climate-controlled (21°C, humidity 65%) vivarium. *Ad libitum* food (LabDiet^®^ JL Rat and Mouse/Irr 6F), water, and environmental enrichment (a combination of Nestlets and either an EcoForage pack or a wood gnawing block and EnviroPak) were provided. Cages were changed 2X/week. Mice were housed in groups of 2–5 until testing and single-housed one week prior to the start of the experiment. Mice were pseudo-randomly assigned (based on body weight) to experimental groups before the start of each experiment. A total of six cohorts were tested in the first experiment, and a total of two cohorts were tested in the second experiment. The first four cohorts were comprised of members of the same sex (all female or all male). Each of the final four cohorts consisted of an equal number of females and males. Experiments took place over a 26-month period, from 2017 to 2020. Mice were not handled by experimenters prior to testing. All experimental protocols were approved by the Vassar College Institutional Animal Care and Use Committee. Disclosure of animal housing, husbandry, and experimental procedures followed principles for transparent reporting (Prager, Bergstrom, Grunberg, & Johnson, 2011) and reproducibility (Prager et al., 2019) in behavioral neuroscience.

### 2.2 Antimicrobial Regimen

The ATMs (2mg/mL neomycin; 2 mg/mL bacitracin; 1.2 μg/mL pimaricin; Sigma-Aldrich, St. Louis, MO) used in this study were administered in tap water to half of all subjects in each cohort for 10 days. Controls received tap water. The ATM cocktail was freshly prepared every second day. In the first experiment, ATMs were started 5 days prior to behavioral testing (Figure 1A); in the second experiment, ATMs were started the day after context recall (Figure 1F). Our cocktail was adapted from prior research showing it is nonabsorbable from the gut, and does not induce intestinal inflammation (Bercik et al., 2011; Odeh, 2013; Verdu et al., 2006; Van der Waaij & Sturm, 1968; Van Der Waaij, Berghuis-de Vries, & Altes, 1974). Intraperitoneal injection of this cocktail has not been shown to induce behavioral changes associated with enteral administration (Bercik et al., 2011), and bacitracin and neomycin do not readily cross the blood brain barrier (Desrochers & Schacht, 1982; Teng & Meleney, 1950).

**Figure 1.**
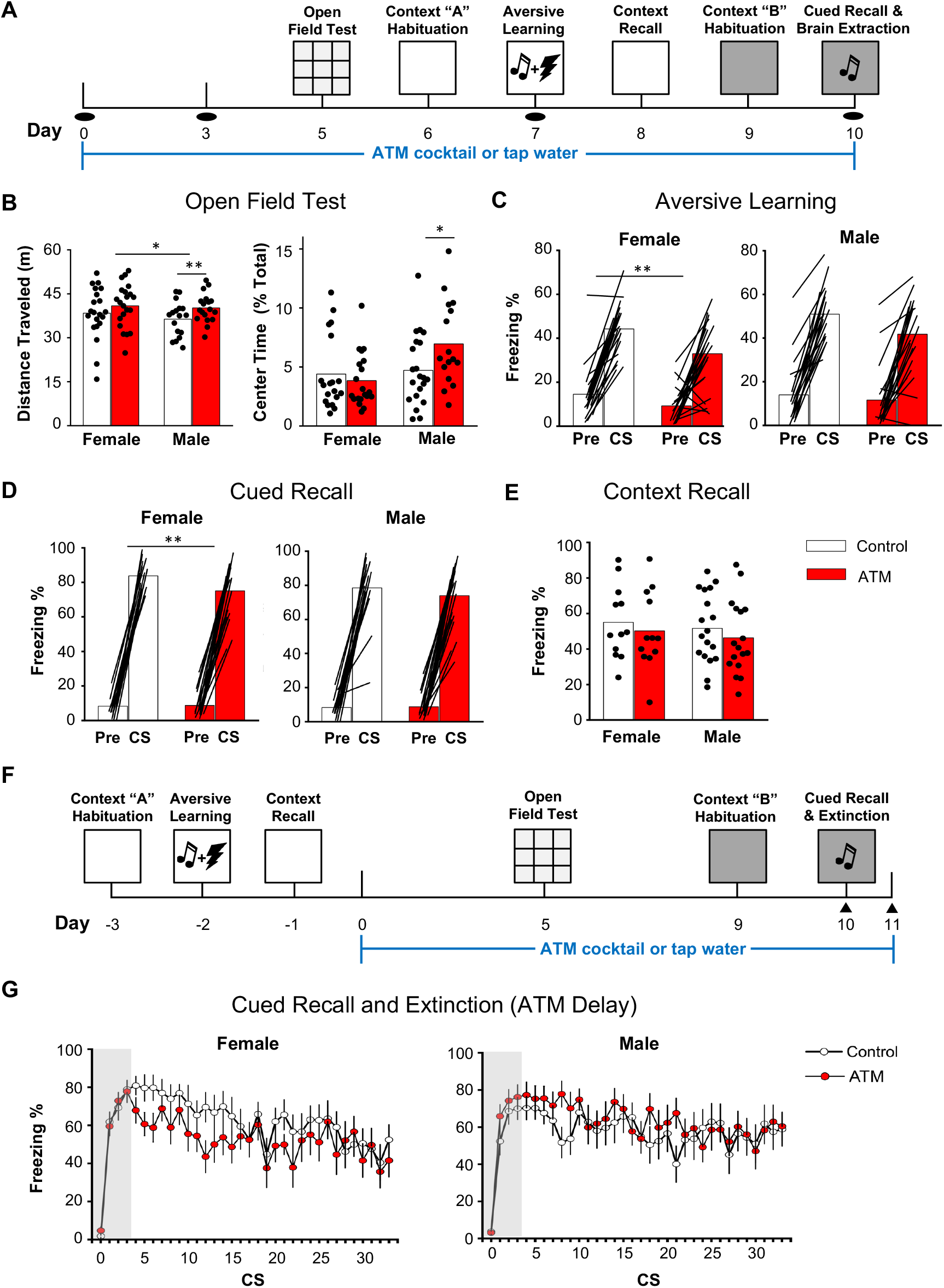
Female C57BL/6N mice are more susceptible than males to gut dysbiosis-driven impairment of cued aversive learning. **(*A*)** Experiment 1 Schematic. Half the subjects were administered ATMs in tap water for 10 days. Controls received tap water. Fecal pellets (black ovals) were collected on Days 0, 3, 7, and 10. **(*B*)** ATMs increased distance traveled in males but not females. Females traveled further than males overall. ATMs increased percentage time spent in the arena’s center in males but not females (n = 18–22/group). **(*C*)** ATMs impaired aversive learning (n = 21–23/group) and **(*D*)** cued recall (n = 19–21/group) in females but not males. **(*E*)** ATMs did not alter context recall in either sex (n = 12–18/group). **(*F*)** Experiment 2 (“ATM Delay”) Schematic. To isolate effects of gut dysbiosis on cued recall, ATMs were started after rather than prior to learning of the aversive cue. Vaginal cytology (black triangles) was performed on Days 10 and 11. **(*G*)** Treatment did not alter cued recall (grey boxed area) or extinction learning (unboxed area) in either sex (n = 12/group). \**P* < 0.05, \*\**P* < 0.01

### 2.3 Animal Health

General health and weight of mice were monitored and recorded daily. Percent body weight change was calculated from baseline (Day 0). In one cohort in the first experiment, ATM dose was dropped to 50% on Day 6 in six female subjects who were approaching > 15% weight loss; dosing was returned to 100% the following day, when body weight increased. All subjects in both experiments displayed normal behaviors and signs of good health (active, alert, well-groomed, clear eyes) throughout ATM administration. Water consumption was assessed only in the first experiment. Water bottles were weighed daily, including prior to placement in or collection from cages. Daily water consumption for each subject was determined by subtracting final from initial bottle weight, then converting that number to mL. To compare water intake across subjects of varying body weights, each subject’s daily water intake (mL) was divided by its daily body weight (g).

### 2.4 Behavioral Testing

Visual depictions of Experiments 1 and 2 are shown in Figure 1. Behavioral testing was conducted during the light cycle. Unless otherwise indicated, all testing equipment was wiped with 70% ethanol before and after each test. Subjects were counterbalanced by treatment and age to minimize potential order effects. The day prior to both fear conditioning (context A) and cued recall (context B), subjects were allowed to freely explore the respective context chamber configuration for 1800 s (“Habituation”).

#### Open Field Test (OFT)

OFT was used to assess baseline locomotor activity (determined by the sum of total distance traveled) and anxiety-like behavior (determined by percentage time spent in the arena’s center, which is considered a more anxiogenic environment than the enshadowed periphery; Bailey and Crawley, 2009). Mice were placed in the bottom-right corner of the novel, dimly-lit rectangular arena (3 lux; 54 cm x 39 cm x 38 cm) and allowed to explore freely for 600 s.

#### Aversive Learning (Context A)

The aversive conditioning chamber measured 18 cm x 18 cm x 45 cm and was enclosed in a sound-attenuating chamber (58 cm x 45 cm x 61 cm; Coulbourn Instruments; Whitehall, PA USA). Programming was delivered via Graphic State software (Coulbourn Instruments). Following a 180 s habituation period, mice were presented with three 20 s auditory cues (CS; 72–75 dB; 5 kHz), each of which co-terminated with a 0.6 mA, 0.5 s foot shock (US). Intertrial intervals (ITIs) varied in length (20, 80, and 60 s). The total training time was 400 s.

#### Context Recall (Context A)

Mice were placed back in the original training context (unmodified commercial chamber) and allowed to explore freely for 1200 s.

#### Auditory Cued Recall (Context B)

Auditory cued recall was conducted in the same chamber as Context A, but the chamber was disguised (hereafter referred to as Context B) to circumvent recall of Context A. Disguises to Context A included modification of habituation conditions (dim lights, cages hand-carried, and white noise, as opposed to bright lights, cages wheeled on cart, and quiet), the aversive conditioning apparatus (1% acetic acid, plastic floor covered in clean bedding, and 2 white stripes added to each plexiglass wall, instead of ethanol, steel bar flooring, and transparent walls), and the experimental protocol (ITI1 and ITI2 were switched, and no foot shocks were delivered). The total test time was 400 s.

#### Auditory Cued Extinction (Context B)

Auditory cued extinction was conducted only in the second experiment, and took place in Context B, as an extension of the cued recall test. Mice were presented with thirty 20s CSs (72–75 dB; 5-kHz). ITIs were created using a random number generator, and varied in length between 5–60 s. The total test time was 1400 s.

### 2.5 Behavioral quantification

Cameras positioned directly above the OFT arena and aversive conditioning chambers recorded video footage. Footage was analyzed using the video tracking system SMART v3.0 (Panlab, Harvard Apparatus, Barcelona, Spain). For OFT, SMART quantified total distance traveled and percentage time spent in the arena’s center (16 cm x 11 cm). For aversive conditioning assays, “freezing” was used as a behavioral measure of a conditioned defensive response. Freezing was defined as immobility, except for respiration, that lasted > 1 s. For each time bin (habituation, CSs, and ITIs), freezing duration was averaged and converted into a percentage. Due to changing technologies within our lab over the course of the experiment, freezing was quantified three different ways. The first four cohorts (Experiment 1) were hand-scored to 95% inter-rater reliability by at least 2 blind experimenters. The following two cohorts (Experiment 1) were quantified using SMART, and the final two cohorts (Experiment 2) were quantified using FreezeFrame (Actimetrics, Wilmette, IL USA). To verify digital tracking software accuracy and comparability of freezing quantification, a subset of the digitally-analyzed footage was hand-scored as before. Comparison of these results with those of the automated tracking software revealed a remarkably high inter-rater reliability (R^2^ = 0.99). Power analyses for determining subject number were carried out using GPower software v.3.1. Power analysis was based on a repeated-measures ANOVA (RM-ANOVA). With a medium effect size (0.50, Cohen’s *d*), high power (0.95) and alpha level of 0.05 required a total of ~100 mice. Therefore, we used approximately 25 mice/group for sufficient power in order to detect a statistically significant effect for behavioral studies.

### 2.6 Behavioral statistics

Prior to analyses, outliers were determined as values greater than or equal to 1.5 times the interquartile range. Analyses were performed using SPSS v. 26 (IBM, Armonk, NY). For all two-way ANOVAs, the between-subject variables were sex and treatment (ATM vs Control). For all within-sex ANOVAs, the between-subjects variable was treatment. Aversive conditioning tests, body weight change, and water consumption were each analyzed using a two-way RM-ANOVA). The dependent variable for each analysis was % freezing, % body weight, and volume liquid consumed, respectively. The within-subjects variable for aversive conditioning analyses was time bin. The within-subjects variable for body weight and water consumption was treatment day. Sphericity was assessed using Mauchly’s Test; if significant, the Greenhouse-Geisser adjustment was applied. Locomotor activity and anxiety-like behavior analyses were conducted using two-way ANOVA. Dependent variables were distance traveled (m) and % time spent in the center of the arena, respectively. Homogeneity of variance tests were conducted using Levene’s test; if significant, the Welch *F*-ratio was used. Because sex differences were hypothesized a priori, all analyses were followed up with within-sex ANOVAs. Bonferonni’s post-hoc tests were conducted as necessary. For relevant comparisons between sex, lower and upper bound 95% confidence intervals (CI) were reported as well as Cohen’s *d* values. We note that our investigation of sex-specific effects was not intended to presuppose rigid binaries, but rather to examine whether sex is associated with variation in our dependent variables (Maney, 2016). All reductions in sample sizes were due to technical error (malfunctioning tracking software and lost footage). All subjects underwent each behavioral test and were excluded from analyses only when data was unavailable. Statistical significance for all data was set at *P* < 0.05. All tests were two-tailed.

### 2.7 Golgi-Cox Staining

The day following auditory cued recall, mice were anesthetized with a ketamine/xylazine cocktail (100:10 mg/mL) and intracardially perfused with saline. The whole brain was submerged in a Golgi-Cox solution composed of 5% w/v mercuric chloride, 5% w/v potassium chromate, and 5% w/v potassium dichromate. Brains were stored in the dark, at room temperature, for 5 days. The solution was refreshed after the second day. Next, brains were transferred to a cryoprotectant (Zaqout & Kaindl, 2016) and stored at 4°C until slicing. Coronal sections 200-*μ*m thick were cut in cryoprotectant using a vibratome (VT1200, Leica Biosystems Inc., Buffalo Grove, IL USA). Sections were transferred to 3% gelatinized slides and left to dry in the dark at room temperature for no more than 2 days. To develop the sections, they were first rinsed in dH20, dehydrated in 50% ethanol, and alkalinized in a 33% ammonia hydroxide solution. Following another rinse in dH20 and immersion in 5% sodium thiosulfate (in the dark), slides were dehydrated in a graded series of ethanol dilutions, cleared using xylenes, and cover slipped using a mounting medium (Permount, Fisher Scientific, Hampton, NH USA). These procedures were adapted from several previous protocols (Bergstrom, Smith, Mollinedo, & McDonald, 2010; Gibb & Kolb, 1998; Zaqout & Kaindl, 2016).

### 2.8 Imaging and 3D dendrite reconstruction

#### Dendrite visualization

Dendrites from Golgi-Cox-stained BLA principal neurons were visualized using brightfield microscopy (Axio Imager M2, Zeiss, Thornwood, NY USA) under a 63X (0.75 N.A.) air objective, and manually reconstructed in 3D using Neurolucida (MBF Biosciences, Williston, VT USA; Figure 2B). All investigators were blind to treatment conditions. Principal neurons in the BLA complex (defined as the lateral, basolateral, and basomedial subnuclei; Figure 2A) were sampled randomly, and every effort was made to sample both hemispheres evenly. Neurons chosen for reconstruction were well-stained (fully impregnated dendritic trees), with unobstructed dendritic arbors that could be followed from soma to terminal tip without interruption. Morphometric analysis was restricted to cells located between Bregma −1.00 mm to −2.30 mm. The BLA was identified by the contour of the two branching fiber tracts that delineate its medial and lateral borders. Principal neurons were differentiated from stellate neurons and interneurons based on morphometric criteria that included spines on later branch orders, an “apical-like” dendritic tree, and biconical dendritic radiation (Bergstrom et al., 2010).

**Figure 2.**
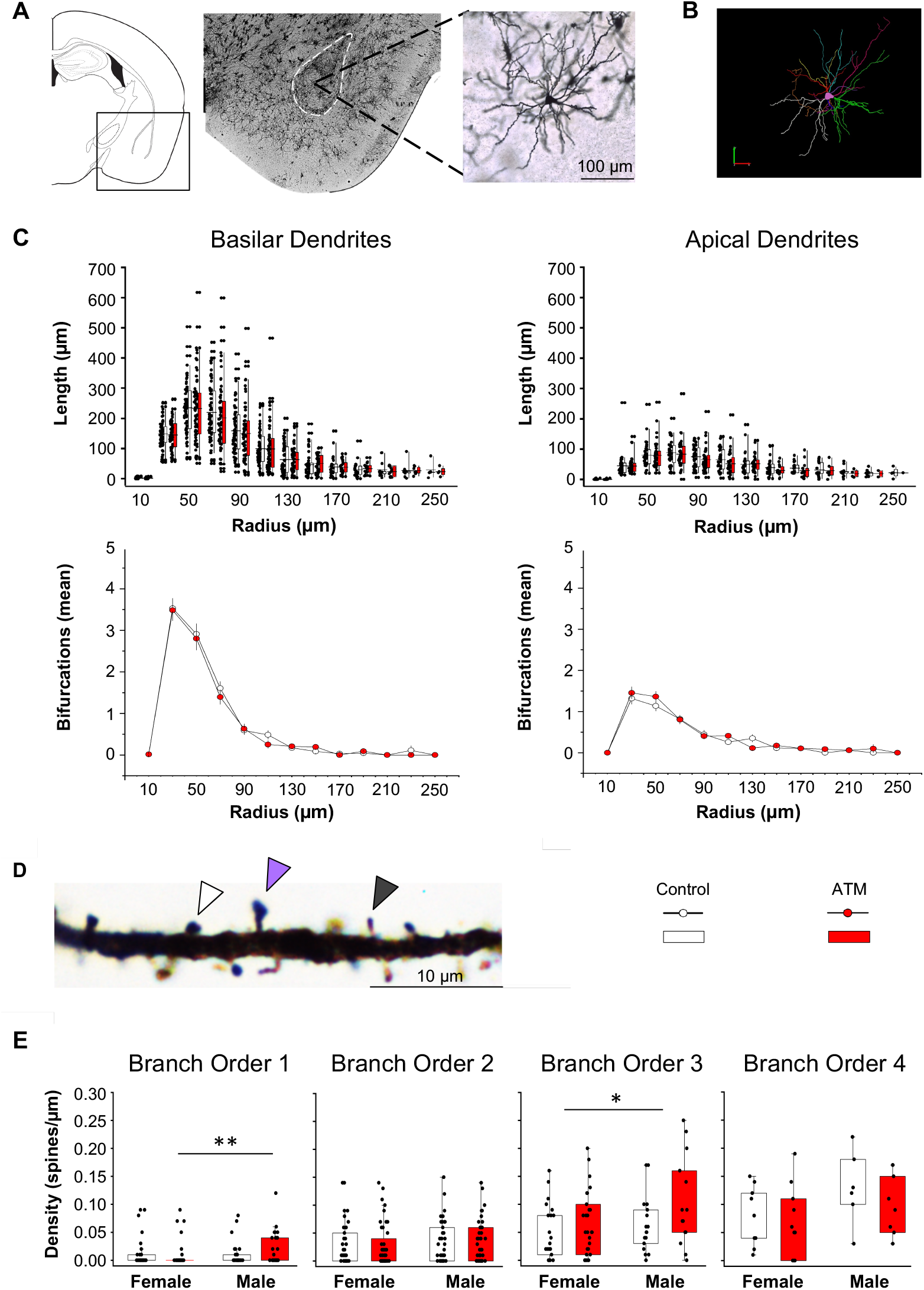
Morphometrics of Golgi-Cox-stained BLA principal neuron dendrites and spines. **(*A*)** The location of the BLA is shown in the atlas (*left*), and enclosed within the dashed lines in the micrograph (20X, *center*). Randomly-sampled principal neurons (40X, *right*) were **(*B*)** reconstructed in 3D using Neurolucida (MBF Biosciences). Mixed model analyses revealed no effects of sex or treatment on **(*C*)** average dendritic lengths or **(*D*)** bifurcations. **(*E*)** Spines (100X) were classified into three morphological types: stubby, mushroom, and thin (white, purple, and grey arrows, respectively). No overall differences were detected between types (data not shown). **(*F*)** No overall differences in spine density were detected, but ATMs exerted an opposing effect between sex at first order branches, with reduced spine density in females relative to males. At third order branches, overall spine density was higher in treated mice than controls (data not shown), and higher in males than females. n = 4–6 mice/group and 35–42 reconstructed cells/group, \**P* < 0.05, \*\**P* < 0.01

#### Dendritic spine visualization

Dendritic spines were visualized under a 100X (1.3 N.A.) oil-immersion objective (Figure 2D). Spines were characterized on a randomly chosen dendritic tree (soma to terminal tip) from a previously-reconstructed neuron. Spines were defined as small (< 3 μm in length) protrusions emanating from the dendritic shaft. Spines were characterized as one of four types: stubby (protrusions that lack a neck), mushroom (neck with a larger head), thin (long, skinny neck and small, bulbous head), or unknown (Yuste, 2010).

### 2.9 Dendrite and dendritic spine morphometric statistics

The sample size for morphometric analyses were based on previous reports (Bergstrom, McDonald, French, & Smith, 2008; Bergstrom, Smith, Mollinedo, & McDonald, 2010). Quantitative morphological measurements were extracted using Neuroexplorer software (MBF Biosciences, Williston, VT). All data were analyzed in R v3.6.1 (R Core Team, 2019). Apical and basilar sholl data were restricted to radii between 30 and 250 μm because there were too few data points to analyze above or below these values. All data were analyzed using generalized linear mixed effects models, with sex and treatment as fixed effects, radius as a random slope, and individual ID as a random intercept (Wilson, Sethi, Lein, & Keil, 2017). The random slope controls for variation in repeated measurements within a single individual (akin to a repeated measures design), while the random intercept controls for multiple cells measured within a single individual. The error distribution for different measures of sholl data was determined by comparing the fit of different possible distributions (i.e., normal, lognormal, Poisson, and negative binomial) with the fitdistr function in the MASS package. Dendritic intersections and overall length were analyzed with a negative binomial distribution using the glmmTMB package (Brooks et al., 2017). Given the large number of zeros in the data (radii without nodes), we first analyzed node data as a binomial variable, investigating the presence/absence of nodes at each radius. We then analyzed radii where nodes were present using a negative binomial distribution to determine if the number of nodes varied by sex or treatment.

Spine data were analyzed in terms of spine density per branch order (spines per μm). Since spine density values were very small (mean = 0.048) and contained a large number of zeros (branches without spines), we analyzed data as generalized linear mixed effects models with a Tweedie distribution using the glmmTMB package. Tweedie distributions (Jørgensen, 1987) are increasingly used in neuroscience (Moshitch & Nelken, 2014), and describe a probability distribution for continuous positive data that includes a large portion of the data at zero, and where the values are generally very small. We began by analyzing the three-way interaction between branch order, sex, and treatment. Random effects included branches and cells nested within individuals. Because this three-way interaction was significant, we then analyzed sex x treatment interactions within each branch order separately, with individual ID as a random effect. Significance was set at P < 0.05 and assessed using likelihood ratio tests of increasingly simplified nested models (Crawley, 2012), essentially equivalent to a two-tailed significance test.

### 2.10 PCR Amplification and Library Preparation

Fecal pellets were collected in two mixed-sex cohorts on Days 0, 3, 7, and 10 of Experiment 1 (Figure 1A). Fecal pellets were not collected during Experiment 2. One pellet was obtained per subject by individually placing mice in a plastic box sterilized with 70% ethanol. Immediately following defecation, sterile tweezers were used to transfer the pellet to a 2.0 mL microfuge tube, which was stored at −80°C. To extract DNA, pellets were heated at 65°C for 10 min, then homogenized and processed using a QIAmp Powerfecal DNA kit (Qiagen). DNA was stored at −20°C.

The 16s rRNA gene amplicon libraries were constructed in a two-stage polymerase chain reaction (PCR) procedure. In the first stage, the V3 and V4 regions of the 16S rRNA genes were amplified with Illumina V3-V4 forward (TCGTCGGCAGCGTCAGATGTGTATAAGAGACA GCCTACGGGNGGCWGCAG) and reverse primers (GTCTCGTGGGCTCGGAGATGTGTATAAGAGA CAGGACTACHVGGGTATCTAATCC). Reactions included 12.5 μL of the 2X KAPA Hifi Hot Start Ready Mix (Kapa Biosystems), 5 μL each of the forward and reverse primers at 1 μM, and 2.5 μL of the template DNA. Thermocycling conditions were as follows: 95°C for 3 min, 95°C for 30 s, 55°C for 30 s, 72°C for 30 s, 72°C for 5 min. Steps 2–4 were repeated 34 times. Gel electrophoresis was used to confirm the presence of 500-600 bp products. Products were quantified using a NanoDrop One (Thermo Scientific, USA).

A QIAquick PCR Purification Kit (Qiagen) was used to purify the original PCR product, then the second-stage PCR was performed to attach Illumina i7 and i5 indexes (Illumina Nextera XT Library Preparation Kit) to individual samples. The reaction contained 12.5 μL of 2X KAPA Hifi Hot Start Ready Mix, 5 μL of the forward and reverse primers at 2μM, and 2.5 μL of the purified 1st stage PCR product. Thermocycling conditions were the same as described above. Steps 2–4 were repeated 8 times. Gel electrophoresis was used to confirm presence of product. A QIAquick PCR Purification Kit was again used to purify second-stage PCR products. After determining concentration using a NanoDrop One, each product was diluted to a concentration of 8 nM prior to combining all samples for sequencing. The final concentration of the combined sample was quantified using Agilent. The sample was sent to Cornell Biotechnology Institute, where the DNA was size-selected using a yBluePippin (Sage Science) before the sample was sequenced on an Illumina MySeq (Illumina pipeline software v2.18) with paired-end 250 base read length.

### 2.11 Gut Microbiome Data Analysis

Paired-end reads were processed and analyzed using QIIME v2018.8 (Bolyen et al., 2019). Reads were trimmed at the 5’ end to remove primer sequences, truncated at the 3’ end to remove low quality ends, and denoised using Dada2 (Callahan et al., 2016). A rooted phylogenetic tree was generated for phylogenetic diversity analysis using the q2-phylogeny plugin, which uses mafft to generate a multiple sequence alignment (Katoh & Standley, 2013) and fasttree to generate the tree (Price, Dehal, & Arkin, 2010). Diversity as a function of sampling depth was investigated using *qiime diversity alpha-rarefaction* with default settings and a maximum depth of 50,000 reads. Three constructs of alpha diversity—Faith’s phylogenetic diversity (Faith, 1992), ASV richness, and species evenness—were calculated using *qiime diversity core-metrics-phylogenetic* with a sampling depth of 10,000 reads per sample. Weighted and unweighted unifrac were calculated using phyloseq in R (McMurdie & Holmes, 2013). Taxonomy was assigned to ASVs using *qiime feature-classifier classify-sklearn* with a SILVA-99 classifier.

All measures of alpha diversity were assessed using linear mixed effects models (LMMs). Analyses were performed in R. All analyses included experimental cohort as a random effect. Thus, our LMMs were analogous to two-way ANOVAs, but they controlled for nonindependence of mice between cohorts. Fixed effects were ATM treatment, sex, and their interaction. Alpha diversity analyses were conducted in two stages. First, we analyzed all data from Day 0 to assess pre-treatment differences. Next, we analyzed data from Days 3, 7, and 10 to assess effects of ATM treatment. Significance was assessed using likelihood ratio tests of increasingly simplified nested models. Model fit was assessed using quantile-quantile plots of residuals.

Whereas alpha diversity assesses complexity within microbial communities, beta diversity assesses complexity between microbial communities (Lozupone & Knight, 2005). Two variants of the UniFrac metric were used to assess phylogenetic distance between communities: unweighted UniFrac, which considers the presence or absence of ASVs, and weighted UniFrac, which considers ASV relative abundance. Both variants were analyzed using permutational multivariate analysis of variance (PERMANOVA), which performed in R using adonis from the package vegan with 999 permutations. Main effects were sex and ATM treatment. Analyses were again conducted in two stages, first for all Day 0 data, then for all treatment days. Significant PERMANOVA results were further analyzed using betadisper from the package vegan. One-way ANOVA was used to determine if results could be explained by differences in dispersion. Principal Coordinates Analysis (PCoA) plots were generated in R to visualize the data. Distances were calculated in phyloseq and ordination was performed using betadisper.

Differential abundance analysis was performed on samples from Days 3, 7, and 10 using the q2-gneiss plugin for QIIME 2 (Morton et al., 2017). ASVs with fewer than 50 sequence counts, or which appeared in under five samples, were removed. Correlation clustering by treatment day was used to define partitions of co-occurring ASVs, which were then log-transformed. Within gniess, a multivariate response linear regression was performed using the ordinary least squares function to identify meaningful covariates (sex, treatment, day) that impacted the model. For subsequent determination of significant balances, gneiss was performed on female and male samples separately using treatment and treatment day for the linear regression. The largest significant balances in both sexes were selected for inspection of taxa populating the balance. Plots were generated in R. Significance for all data was set at *P* < 0.05.

### 2.12 Vaginal Cytology and Histology

Because the influence of ovarian hormones on aversive extinction (but not cued aversive learning or recall) are well-established (Milad et al., 2009; Velasco, Florido, Milad, & Andero, 2019), we conducted a preliminary investigation of potential ATM x estrous stage interactions during the second experiment. Vaginal cells were collected on Day 10, after the cued recall/extinction test, then again on Day 11 to confirm estrous stage identification accuracy. Each mouse was held by its tail, paws resting on the cage lid, while 2 mL of sterile water were pipetted in and out of the vaginal opening. The pipette did not penetrate the orifice. Water containing cells was released onto a glass slide and left to dry. Once dry, estrous smears were submerged in a 0.1% crystal violet solution for 1 min, then exposed to two 1 min rinses in distilled water prior to coverslipping. Slides were examined using brightfield microscopy at magnifications of 40X and 100X. Cornified squamous epithelial cells, leukocytes, and nucleated epithelial cells were identified, and their relative proportions were used to determine estrous stage as previously described (Byers, Wiles, Dunn, & Taft, 2012).

### 2.13 Estrous Stage Statistics

We created a binary analysis by combining diestrus and proestrus (stages in which levels of ovarian hormones are relatively high) as well as estrus and metestrus (stages in which levels of ovarian hormones are relatively low). To assess effects of ovarian hormones on cued recall and extinction, the first 5 CSs and last 5 CSs were collapsed and compared using a two-way RM-ANOVA. Because treatment differences were hypothesized a priori, one-way ANOVAs within estrus and diestrus were also performed. Analyses were performed as described above in the behavioral statistics section.

## 3 RESULTS

### Experiment 1: ATMs on Aversive Conditioning

#### 3.1 Open Field Test (OFT)

Two-way ANOVA on total distance traveled (m) revealed main effects of sex (*F*_1, 76_ = 5.1; *P* = 0.03, *d* = 0.61) and treatment (*F*_1, 76_ = 7.0; *P* = 0.01, *d* = 0.74), with no interaction (Figure 1B). Females (n = 41, mean = 40.2 ± 1.2) traveled further than males (n = 39, mean = 36.3 ± 1.3), and treated mice (n = 39, mean = 40.5 ± 1.4) traveled further than controls (n = 41, mean = 35.9 ± 1.2). ATMs increased distance traveled in males (treated: n = 18, 95% CI [37.7, 42.7]; controls: n = 21, 95% CI [27.9, 36.9]) (Welch *F*_1, 30.4_ = 9.9, *P* = 0.004, *d* = 0.84), but did not affect distance traveled in females.

Two-way ANOVA on percentage time spent in the field’s center revealed a sex x treatment interaction (*F*_1, 75_ = 4.4; *P* = 0.039; Figure 1B). Within males, percentage center time was increased in treated subjects (n = 18, 95% CI [5.7, 10.5]) relative to controls (n = 21, 95% CI [3.3, 6.1]) (Welch *F*_1, 27.7_ = 6.7, *P* = 0.015, *d* = 0.74). ATMs did not affect percentage center time in females. Together, these data suggest that perturbation of the gut microbiome increased locomotor activity and reduced anxiety-like behavior in males but not females.

#### 3.2 Aversive Learning

Two-way RM-ANOVA revealed effects of sex (*F*_1,82_ = 9.95, *P* = 0.002, *d* = 0.87) and treatment (*F*_1,82_ = 11.1; *P* < 0.001, *d* = 0.91; Figure 1C), with no interaction. Within females, treatment reduced freezing in ATM mice (n = 23, 95% CI [32.5, 37.9]) relative to controls (n = 22, 95% CI [36.6, 48.9]) (*F*_1,43_ = 6.3; *P* = 0.016, *d* = 0.79), suggesting impaired learning of the CS/US association. Treatment did not affect freezing in males. There was no difference in freezing between female and male controls. Importantly, there were also no pre-CS/US differences in freezing. These data suggest that females are more susceptible than males to gut dysbiosis-related modulation of aversive learning.

#### 3.3 Context Recall

Two-way RM-ANOVA did not detect a significant effect of sex or treatment, or any interactions (Figure 1D).

#### 3.4 Cued Recall

Two-way RM-ANOVA revealed an effect of treatment (*F*_1,70_ = 9.2; *P* = 0.003, *d* = 0.82), such that treated mice showed less freezing than controls (Figure 1E). There was no effect sex, or interaction of sex x treatment. Within females, treated mice froze less (n = 19, 95% CI [54.9, 75.7]) than controls (n = 19, 95% CI [73.4, 86.6]) (*F*_1,36_ = 6.3, *P* = 0.017, *d* = 0.68), suggesting impaired memory retrieval of the CS/US association. Treatment did not affect freezing in males, and there were no differences in freezing between female and male controls. There were also no differences in pre-CS freezing. These data suggest that females are more susceptible than males to gut dysbiosis-related modulation of cued aversive recall.

#### 3.5 Dendritic and Spine Morphology of BLA Principal Neurons

Effects of gut dysbiosis on aversive conditioning led us to investigate potential changes in BLA principal neuron morphology. Generalized linear mixed effects modeling did not reveal effects of ATMs on apical or basilar dendritic structure (Figure 2C), but effects were detected for dendritic spines. Analysis of spine density revealed a threeway interaction between branch order, sex, and treatment, indicating that spine density was affected by treatment and sex, but effects differed across branch order (*χ*^2^ = 9.9, *P* = 0.019; Figure 2E). Branch order analyses revealed effects at 1st and 3rd order dendrites. For the 1st order tree, there was a sex x treatment interaction (*χ*^2^ = 4.4, *P* = 0.035) which indicated that ATMs decreased spine density in females relative to males (post-hoc comparison; *χ*^2^ = 5.1, *P* = 0.025). There were no differences between female and male controls. At the 3rd order tree, there were effects of sex (*χ*^2^ = 4.0, *P* = 0.046) and treatment (*χ*^2^ = 4.0, *P* = 0.046), but no interaction. Males had a higher density of spines than females, and treated mice had a higher density of spines than controls. No effects were detected at 2nd and 4th order trees.

Analysis of spine morphology indicated that over 50% of spines on 1st order branches in treated mice were “thin” type spines, compared with fewer than 10% in controls. There were no differences between groups across “stubby” and “mushroom” spine types.

#### 3.6 16S rRNA Sequencing

We obtained over 14 million reads from 53 samples, with > 5,000 reads each. After denoising with Dada2, there were over 10 million reads and 52 samples with > 5,000 reads. Calculation of the Shannon index diversity metric on rarefied samples revealed that 5,000 reads were sufficient to capture sample diversity, although observed ASVs and PD continued to rise steadily until 20,000 reads (data not shown). All further analyses were performed using 52 samples containing at least 5,000 reads to balance sequencing depth and sample retention.

#### 3.7 Gut Microbiome Structure

LMMs revealed no differences between groups for any measure of alpha diversity (PD, evenness, and ASV richness) on Day 0 (Figure 3A). These data indicate that, prior to ATM administration, gut microbiomes were similar between females and males as well as between mice assigned to control and treatment groups. Initial analyses of alpha diversity measures for treatment days (Days 3, 7, and 10) revealed no effect of day, so these days were collapsed. LMMs revealed a main effect of treatment on PD (χ^2^ = 10.5, *P* = 0.001), evenness (χ^2^ = 23.8, *P* < 0.001), and richness (χ^2^ = 21.8, *P* < 0.001), with ATM mice showing reductions in all measures relative to controls. There was also a main effect of sex on PD (χ^2^ = 9.6, *P* = 0.002), evenness (χ^2^ = 4.8, *P* = 0.03), and richness (χ^2^ = 4.9, *P* = 0.03). The interaction of sex x treatment did not affect evenness or richness, but it affected PD (χ^2^ = 3.9, *P* = 0.047) such that PD decreased more in ATM females than ATM males. Together, these results indicate that female mice are more susceptible to effects of ATM treatment on PD than males, but otherwise ATMs generally impact gut microbiomes similarly across sex.

**Figure 3.**
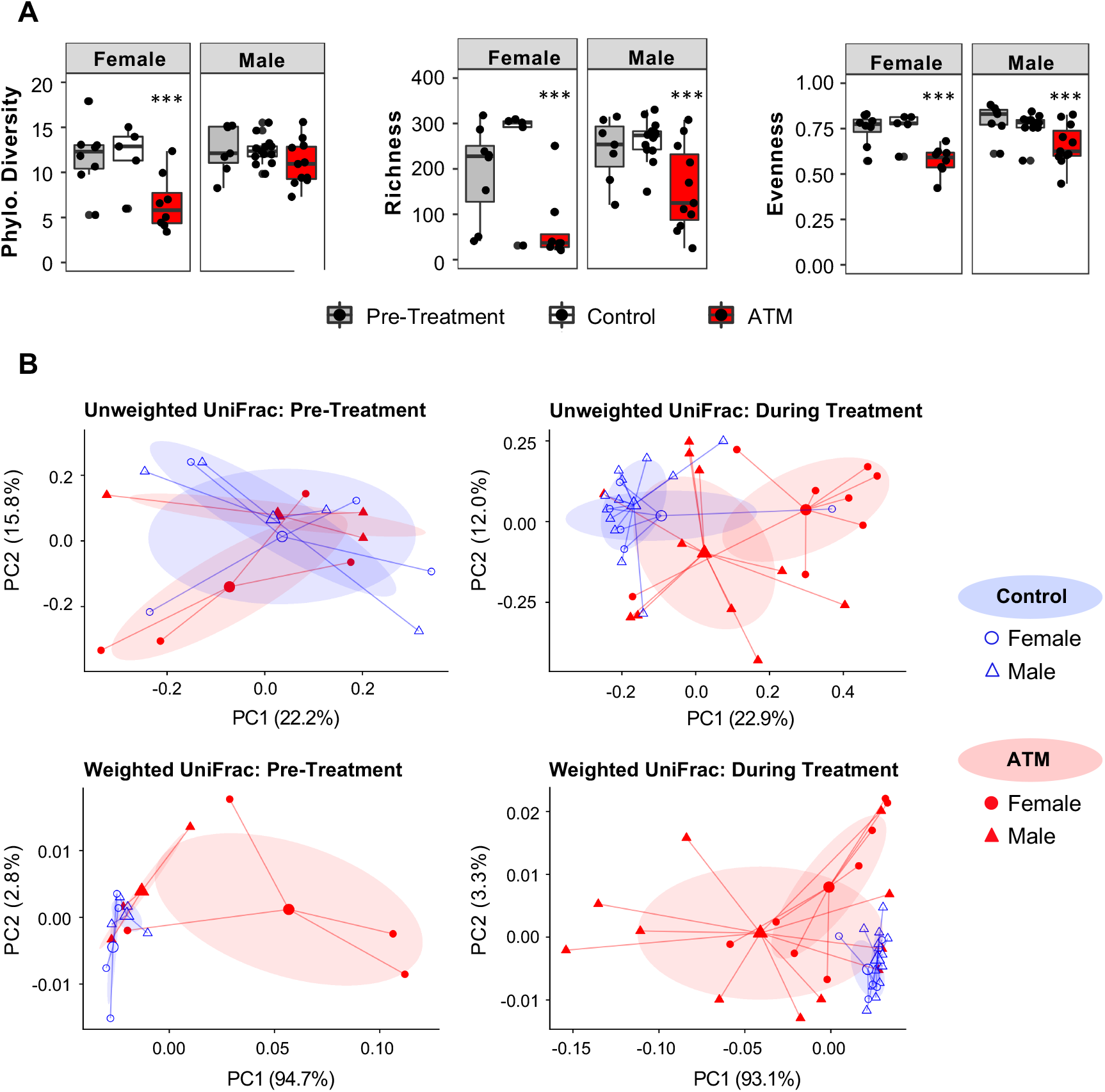
ATMs induce gut dysbiosis in both sexes, with stronger effects in females. **(*A*)** ATMs reduced ASV richness and evenness in both sexes and reduced phylogenetic diversity only in females. These effects did not differ across Days 3, 7, and 10. **(*B*)** Principal coordinate (PC) plots of unweighted and weighted UniFrac distances revealed that microbiomes of ATM mice became increasingly dissimilar after the start of ATM administration, indicating that treatment altered community structure. PERMANOVAs revealed no effects of sex, except for unweighted UniFrac during treatment (*P* < 0.01). This effect was not due to differences in dispersion, and indicates that ATMs differentially impacted rare species between sex. Large shapes represent centroids, small shapes represent samples. Shaded ovals indicate one standard deviation from the centroid. n = 5–13 samples/group, \*\*\**P* < 0.001

Beta diversity was assessed using both unweighted (qualitative) and weighted (quantitative) variants of the UniFrac metric. Prior to treatment (Day 0), PERMANOVA of unweighted UniFrac distances revealed no differences between groups (Figure 3B). PERMANOVA of weighted UniFrac distances revealed an effect of treatment (pseudo-*F* = 6.6, *P* = 0.006) which was partly due to differences in dispersion (*F*_1,13_ = 7.4, n = 15, *P* = 0.017). Visualization of the PCoA plots revealed a likely difference in centroids between ATM and control groups of both sexes.

Initial analyses of unweighted and weighted UniFrac distances revealed no effect of ATMs across treatment day (Days 3, 7, and 10) in either sex, so treatment days were collapsed. This finding suggests that ATMs exerted relatively rapid and persistent changes in gut microbial communities. PERMANOVA of unweighted UniFrac distances revealed an effect of both treatment (pseudo-*F* = 3.57, *P* = 0.001) and sex (pseudo-*F* = 2.61, *P* = 0.004), with no interaction. Differences in dispersion partially accounted for the effect on treatment (*F*_1,35_= 8.12, n = 37, *P* = 0.008), but not sex. These findings suggest that differences between females and males were due entirely to varying positions in centroids. PERMANOVA of weighted UniFrac distances revealed an effect of treatment (pseudo-*F* = 13.1, *P* = 0.001), which was due at least in part to differences in dispersion (*F*_1,35_ = 20.29, n = 37, *P* < 0.001). There was no effect of sex. Visualization of the PCoA plots also suggested a difference in centroids of both sexes between ATM and control groups. Together, these results confirm that ATMs were ingested and induced states of gut dysbiosis in both females and males. Because an effect of sex was detected only when considering ASV presence or absence, our data further suggest that differences between ATM females and ATM males occurred only in rare ASVs (taxa in relatively low abundances).

We next sought to characterize gut microbiome taxonomic shifts (Figure 4A). In both sexes, microbiomes of controls were dominated by Bacteroidetes and Clostridia. ATM mice of both sexes showed reductions in the relative abundance of Clostridia, and increases in Verrucomicrobia and Gammaproteobacteria.

**Figure 4.**
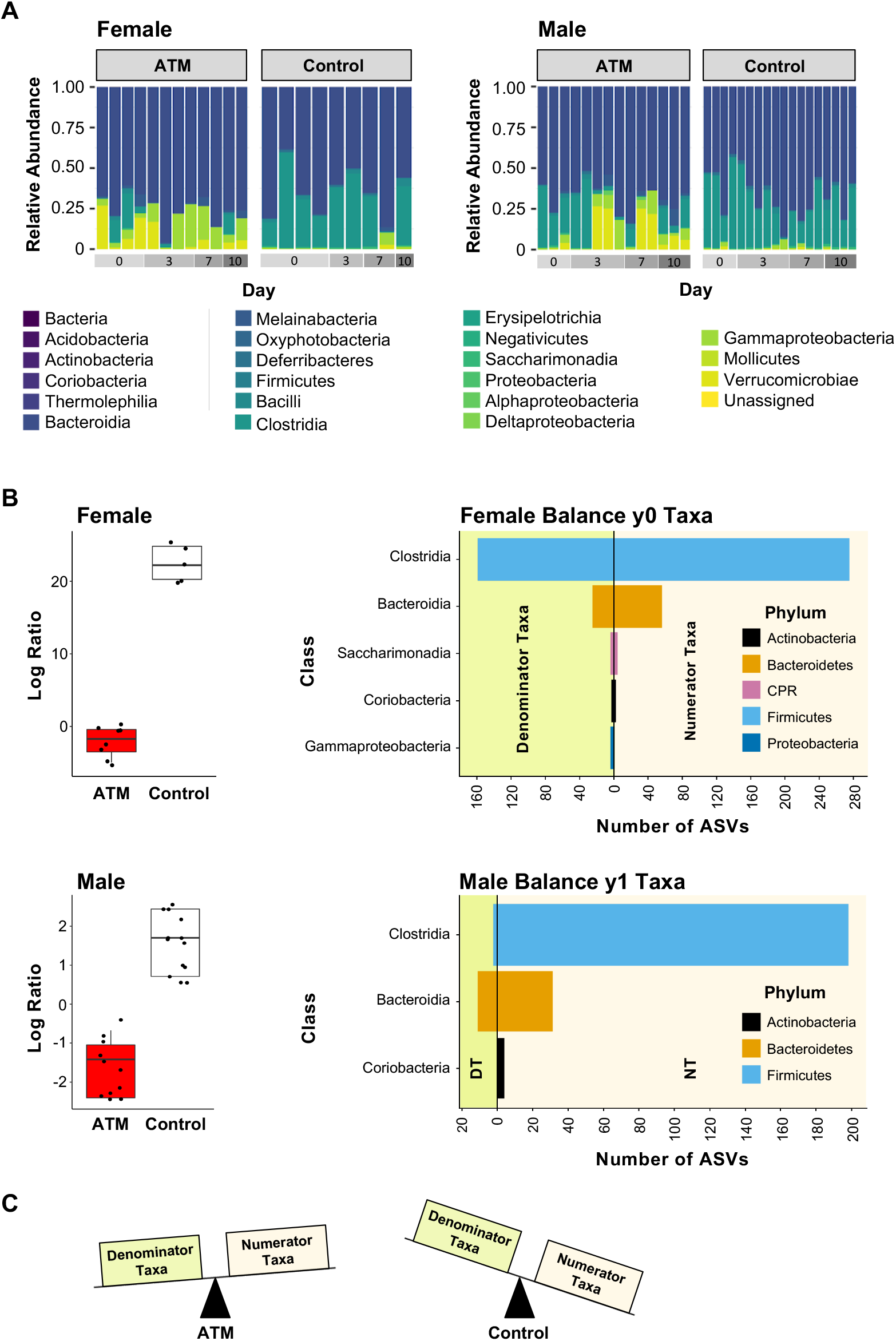
ATMs alter gut microbiome taxonomic structure. **(*A*)** Relative abundance of major phyla across the experiment. **(*B*)** Gneiss was used to calculate log ratios of numerator:denominator taxa in both sexes to reveal differences in microbial sub-communities (called balances) between ATM mice and controls during treatment days. Shown for each sex is the log ratio of the largest significant balance (*left*) and the taxa comprising the balance (*right*). Taxa represented by 4 or less ASVs were excluded for clarity. Note that these values represent the taxonomic composition of the sub-communities, not relative abundance of the ASVs. Both sexes showed a downward shift in the log ratio for ATM mice relative to controls, **(*C*)** visually depicted by the two scales. Given that we expect ATMs to deplete microbial populations, this shift most likely indicates that treatment either decreased relative abundance of numerator taxa, or it decreased relative abundance of both numerator and denominator taxa, with a larger decrease in the former. males: n = 25/group, females: n = 13/group

Differential abundance analysis was performed using gneiss, which uses the concept of balance trees to determine shifts in microbial subcommunities. Linear modelling identified treatment as the main covariate explaining variation in the communities (10.9%). Sex explained 5.4% and treatment day explained 3.0%. Subsequent linear regressions conducted within sex revealed that treatment explained 22.0% of the variation among females and 11.4% of the variation among males. Differences between log ratios of the abundances of bacterial sub-communities (the balances) were found in ATM mice of both sexes relative to controls. Log ratios of the significant balance (*P* < 0.001) containing the largest number of taxa are shown in Figure 4B. A log ratio of 0 would indicate that the relative abundances of taxa in the numerator and denominator were equal. ATM mice of both sexes showed a downward shift in the log ratio of this balance, indicating a transition in dominance away from numerator taxa. This could indicate any of the following possibilities: increased relative abundance of denominator taxa; decreased relative abundance of numerator taxa; a combination of these two; decreased numerator and denominator taxa, with a greater decrease in numerator taxa. Given that we expected ATMs to deplete microbial diversity, the second and last possibilities are the most likely explanations for our data. In both females and males, the taxa in the sub-communities that accounted for the largest differences between ATM and control microbiomes were almost entirely Clostridia and Bacteroidetes (Figure 4B).

#### 3.8 Body Weight

Two-way RM-ANOVA revealed effects of sex (*F*_1,77_ = 16.9, *P* < .001, *d* = .98) and treatment (*F*_1,77_ = 475.5, control n = 47 and ATM n = 47, *P* < .001, *d* = 1; Figure S1A), with no interaction. Both sexes (female n = 46 and male n = 48) showed an equivalent initial drop in body weight in response to ATMs (Days 1–6). Treated females showed a slight rebound in body weight across Days 7–10 (Bonferroni-corrected ANOVAs, *P* < .01).

#### 3.9 Water Consumption

Two-way RM-ANOVA revealed an effect of treatment x day (*F*_8,160_ = 3.2, *P* = 0.002, *d* = 0.97; Figure S2). Bonferroni-corrected follow-up ANOVAs revealed no differences at any particular time point. Within females (n = 46), there was a treatment x day interaction (*F*_8,96_ = 4.7, *P* < 0.001, *d* = 0.99) that showed reduced consumption in ATM mice relative to controls on Days 1–4 and 10 (Bonferroni-corrected ANOVAs, *P* < .01). No differences were detected within males (n = 48).

### Experiment 2: ATMs on Cued Aversive Recall and Extinction (“ATM Delay”)

#### 3.10 Open Field Test

Two-way ANOVA on total distance traveled detected no significant effects, indicating that locomotor activity was not affected by treatment, and did not differ between sex. Two-way ANOVA on percentage center time revealed an increase in the ATM group relative to controls (*F*_1, 44_ = 4.8; *P* = 0.03, *d* = 0.57), suggesting that gut dysbiosis reduced anxiety-like behavior. This effect largely replicated the previous experiment. Within-sex analyses were also similar to the first experiment; percentage center time did not differ between females groups, but within males, there was a trend towards significance between ATM mice and controls (*P* = 0.08; Figure S4A).

#### 3.11 Aversive Learning and Context Recall

Two-way RM-ANOVA detected no significant effects in either aversive learning or context recall (Figure S4B,C), indicating that there were no baseline differences between sex or treatment groups prior to ATM administration.

#### 3.12 Cued Expression (Recall and Extinction)

Cued recall and extinction learning were each assessed using two-way RM-ANOVA. No sex, treatment, or interaction effects were detected (Figure 1G; S4D), indicating that gut dysbiosis does not alter mechanisms associated with either cued memory retrieval or extinction learning.

#### 3.13 Estrous Cycle Stage and Cued Extinction

Mixed RM-ANOVA revealed a main effect of estrous stage on aversive extinction (*F*_1, 20_ = 11.6; *P* = 0.003, n = 6 in estrus/metestrus and n = 6 in diestrus/proestrus for both ATM mice as well as controls). Relative to mice in estrus/metestrus, mice in diestrus/proestrus exhibited less freezing during late extinction only (CSs 29-33; (*F*_1, 20_ = 4.97, *P* = 0.03; Figure S3B)), suggesting that high levels of ovarian hormones augment aversive extinction. One-way ANOVA within subjects in diestrus/metestrus revealed a decrease in freezing in ATM mice relative to controls for CSs 1-5 (*F*_1, 10_ = 8.9; *P* = 0.013), but not CSs 29-33 (Figure S3C). These results suggest that gut dysbiosis may interact with estrous stage to attenuate retrieval of an aversive CS. No effects were detected for oneway ANOVAs within subjects in estrus/metestrus, suggesting that this interaction effect occurs only when ovarian hormone levels are relatively high.

#### 3.14 Body Weight

Two-way RM-ANOVA revealed main effects of sex (*F*_13, 260_ = 2.2, female n = 24 and male n = 24, *P* =.01, *d* = 0.96) and treatment (*F*_13, 260_ = 9.9, control n = 24 and ATM n = 24, *P* < 0.001, *d* = 1.0), as well as a sex x treatment interaction (*F*_13, 260_ = 1.9, *P* < 0.02, *d* = 0.93; Figure S1B). Differences in body weight emerged on Day 4 of ATM treatment (Bonferroni-corrected ANOVAs, *P* < 0.05), with greater percent weight loss in the ATM treatment group relative to controls. In females, there was a rebound in body weight beginning on Day 7 of ATM treatment. Both of these results replicate those of experiment 1 (Figure S1A).

## 4 DISCUSSION

The past two decades have witnessed remarkable strides in understanding the role that gut microbiota play in human health, including in trauma and anxiety-related disorders. A growing body of research has given rise to a framework for how gut microbiota influence aversive conditioning and associated neurobiological mechanisms of plasticity (Cowan et al., 2018), but our understanding of these processes remains nascent. As is the case in much neuroscience research, sex differences remain largely unexplored (Shansky & Woolley, 2016). In this study, we investigated effects of ATM treatment on gut microbiome structure, aversive conditioning, anxiety-like behavior, and BLA principal neuron dendritic architecture in female and male B6 mice. We found wide-ranging and sex-specific effects of ATMs on gut bacterial communities, auditory cued aversive learning and memory recall, anxiety-like behavior, locomotor activity, and BLA dendritic spine patterning. No differences in context recall were detected. We also conducted a second experiment to determine whether gut dysbiosis specifically impacts aversive memory retrieval. When ATMs were administered after memory consolidation, treated subjects showed no behavioral differences relative to controls, indicating that perturbations in gut microbiota uniquely affect aversive learning mechanisms.

### 4.1 Gut microbiome and aversive conditioning

In the first experiment, no differences in conditioned freezing were observed in ATM males. This is consistent with a recent study which found no differences in aversive learning and cued recall in males treated with a different ATM cocktail, or who were of GF or gnotobiotic status (Chu et al., 2019). In contrast, ATM females showed reduced freezing relative to controls during both aversive acquisition and cued recall, suggesting impaired learning of the aversive cue. This finding, to our best knowledge, is novel, and suggests a female-specific susceptibility to gut microbial modulation of aversive conditioning using an ATM cocktail. Because no differences in cued recall were detected in either sex in the ATM Delay experiment, this female-specific susceptibility appears to be due to alterations in learning performance or memory consolidation, as opposed to amygdala-dependent retrieval mechanisms.

Other studies have reported that microbiota can modulate aversive conditioning in males. Two studies in B6 males—one examining GF status (Hoban et al., 2018) and another examining effects of a fecal transplant from mice fed a high-fat diet (Bruce-Keller et al., 2015)—found impairments in cued aversive recall, indicating disrupted memory consolidation or retrieval, as opposed to learning performance. Probiotic studies in BALB/c males provide additional evidence and add nuance: *Bifidobacterium breve* and *B. longum* were found to augment aversive learning (Savignac, Kiely, Dinan, & Cryan, 2014), and both the latter strain and *Lactobacillus rhamnosus* were found to augment cued recall (Bravo et al., 2011). Together, these data suggest that different microbial strains, both individually and synergistically, can exert diverse effects on aversive learning and memory, and that these effects can differ between sex.

Whereas cued aversive recall depends on the amygdala, context recall depends on the hippocampus (Chaaya et al., 2019). In both experiments, we found no effect of ATMs on hippocampal-dependent memory, either in the context recall test or the pre-CS (habituation) period of cued recall. These results are consistent with previous work in male mice in GF (Hoban et al., 2018) and fecal transplant studies (Bruce-Keller et al., 2015). However, several GF studies suggest that microbiota modulate hippocampal structure and function (Clarke et al., 2013; Heijtz et al., 2011; Luczynski et al., 2016). Given that GF animals show developmental abnormalities across nearly all organ systems, including alterations in amygdala and immune function (Al-Asmakh & Zadjali, 2015; Mayer, Savidge, & Shulman, 2014; O’Mahony, Clarke, Gibney, Dinan, & Cryan, 2011; Stilling et al., 2015), the question arises whether gut microbiota influence the hippocampus only within a limited developmental window, prior to adulthood. Models of gut dysbiosis in adult mice do not support this possibility, and MGB studies in normal adult males—including one which used the same ATM cocktail (Bercik et al., 2011)—do not support this hypothesis (Bravo et al., 2011; Provensi et al., 2019; Savignac et al., 2015).

Extinction was conducted only in the second experiment. We observed no effects of ATMs, or differences between sex. Our data contrast with previous reports that microbiota modulate extinction in male rodents (Bravo et al., 2011; Chu et al., 2019; Fox et al., 2017; Hoban et al., 2018; Savignac et al., 2015). However, in agreement with previous work that high levels of ovarian hormones facilitate aversive extinction (Milad et al., 2009; Velasco, Florido, Milad, & Andero, 2019), we found that female ATM mice and controls in diestrus/proestrus, but not estrus/metestrus, showed reduced freezing during the final 5 presentations of cued expression. We also observed that ATMs interacted with females in diestrus/proestrus to attenuate the first 5 CSs of cued expression, suggesting a distinct effect of gut dysbiosis on cued aversive recall when levels of ovarian hormones are high. We note that our second experiment was smaller in scale than the first and should be replicated in larger samples.

### 4.2 Gut microbiome and locomotor activity

Microbial modulation of locomotor activity has been inconsistently reported. In the first experiment, we found that ATMs increased distance traveled in the novel open field in male but not female mice. This supports previous work that GF males show enhanced OFT activity, a phenotype which is reversible by colonization with indigenous gut microbiota, or particular strains of bacteria (Heijtz et al., 2011; Nishino et al., 2013). The sex differences we observed are also consistent with a study in the same strain: B6 males but not females that are fed a high-fat diet and exposed to chronic mild unpredictable stress show reduced locomotor activity (Bridgewater et al., 2017). Together, these data suggest that males are more susceptible to microbial modulation of locomotor activity than females, and this modulation is not developmentally primed.

In contrast, we found no evidence of differences in locomotor activity in the second experiment. A lack of difference is a more widely reported finding across rodent strains and methods of gut microbiome manipulation (Bruce-Keller et al., 2015; Burokas et al., 2017; De Palma et al., 2017; Neufeld, K. M., Kang, Bienenstock, & Foster, 2011a; Savignac et al., 2014; Zheng et al., 2016). This body of work includes a study using female and male B6 mice (Davis et al., 2017) as well as a study using the same ATM cocktail (Bercik et al., 2011), although the latter observed increased activity in non-OFT behavioral assays.

In the first experiment only, we found that females traveled further than males overall. This finding is consistent with a large body of literature across rodent models, including B6 mice (Bridgewater et al., 2017; Foldi, Eyles, McGrath, & Burne, 2011; McCormick, Robarts, Kopeikina, & Kelsey, 2005; Ramos, Correia, Izídio, & Brüske, 2003). However, sex differences in murine locomotor activity are not ubiquitously reported (Davis et al., 2017; De Palma et al., 2017; Leclercq et al., 2017), and no differences were detected in our second experiment. We questioned whether impairments in cued learning and recall in the first experiment could be due to sex-specific locomotor baseline differences. This possibility is unlikely, as locomotor activity between treated and control females was statistically equivalent.

### 4.3 Gut microbiome and anxiety-like behavior

In the first experiment, ATMs increased OFT percentage center time in males, but not females, indicating that altering the gut microbiome induced a sex-specific reduction in anxiety-like behavior. Results from the second experiment, while limited in comparability due to exposure to the fear learning paradigm, also trended towards increased center time in ATM males, and detected no difference between female treatment groups. This anxiolytic phenotype is consistent with numerous studies using a variety of behavioral tests (including OFT) in GF and ATM-treated male mice (Bercik et al., 2011; Clarke et al., 2013; Desbonnet et al., 2015; Heijtz et al., 2011; Zheng et al., 2016), suggesting that dysbiosis or absence of resident microbiota decreases anxiety-like behavior. Prebiotics and probiotics have also been shown to reduce anxiety-like behavior in male mice (Bravo et al., 2011; Burokas et al., 2017; Savignac et al., 2014). Conversely, gut dysbiosis induced by fecal transplants from mice fed a high fat diet (Bridgewater et al., 2017; Bruce-Keller et al., 2015), or from human patients with major depressive disorder (Kelly et al., 2016; Zheng et al., 2016), has been shown to increase anxiety-like behavior in male rodents. Together, these data suggest that the mechanisms underlying microbial modulation of trait anxiety are diverse, and specific to particular bacterial strains and interacting microbial communities.

Few studies have examined whether gut microbiota similarly modulate anxiety-like behavior in females. Consistent with our data, a small but growing pool of evidence suggests that adult females are less susceptible to MGB modulation of anxiety-like behavior than adult males. One study reported an anxiolytic phenotype in female GF Swiss Webster mice, but colonization with resident microbiota in adulthood did not reverse this phenotype (Neufeld, Karen-Anne M., Kang, Bienenstock, & Foster, 2011b) as it did in males of the same strain (Clarke et al., 2013). Since the latter study conventionalized subjects at 3-weeks of age, it is possible that this discrepancy reflects a critical developmental window, rather than a sexspecific effect. However, two studies in B6 mice also support a male-specific vulnerability to MGB modulation of anxiety-like behavior in adulthood: dietary supplementation with docosahexaenoic acid (a long-chain omega-3 polyunsaturated fatty acid) reduced anxiety-like behavior in socially isolated males, but not females (Davis et al., 2017), and chronic consumption of a high-fat diet increased anxiety-like behavior in males, but not females (Bridgewater et al., 2017). While our findings add to mounting evidence for sex-dependent influences of gut microbiota on anxiety-like behavior in adult mice, anxiety is a multidimensional trait, and future research would benefit from systematic examination using multiple behavioral tests. We also note that, because neuroscience research has historically demonstrated sex bias (the tendency to study only male subjects), indices of anxiety which are well-defined in males may be differentially expressed in females; there is a need to investigate potential sex-specific behavioral response profiles (Shansky, 2016).

### 4.4 Gut microbiome and BLA principal neuron dendritic and spine morphology

The amygdala is a well-established network node in the encoding, storage, and retrieval of aversive CS/US associations, and affiliated defensive behavioral responses (Bergstrom, 2016). The BLA plays a particularly crucial role in aversive conditioning, and also influences innate anxiety-like behavior (Cowan et al., 2018; Felix-Ortiz et al., 2013; Simpson, Neria, Lewis-Fernández, & Schneier, 2010). We found that ATMs exerted sexspecific effects on BLA principal neuron dendritic spine patterning, with females showing decreased spine density relative to males on first order branches only. Interestingly, the majority of these spines were “thin” type, a morphology sometimes called the “learning spine” due to its rapid emergence and retraction in response to synaptic activity as well as its potential to grow into a more stable type associated with memory storage (Bourne & Harris, 2007). Our data are consistent with a recent study indicating that BLA principal neurons in GF males have increased spine density across morphological types (Luczynski et al., 2016). The same study also observed elongated BLA principal neuron dendrites, with no differences in bifurcations. We did not detect changes in either of these measures, but it would be useful to explore whether a longer ATM regimen might yield different results. Although our study was similar in length to those using the same cocktail (Bercik et al., 2011; Odeh, 2013; Verdu et al., 2006), other ATM studies employ far longer regimens (Hoban et al., 2016; Kelly et al., 2016). Overall, our data add to growing evidence that gut microbiota influence the amygdala (Bercik et al., 2011; Hoban et al., 2016; 2017; 2018; Neufeld et al., 2011a), particularly the BLA (Bravo et al., 2011; Chu et al., 2019; Heijtz et al., 2011; Luczynski et al., 2016), and—because dendritic structure reflects function (Chklovskii et al., 2004)—suggest that morphological alterations in BLA principal neuron spine density may relate to changes in aversive memory processing and anxiety-related behavior.

### 4.5 ATM-induced shifts in gut microbiome composition

We found that ATMs reduced phylogenetic diversity only in females, and perturbed ASV richness and evenness to a similar degree in both sexes. UniFrac analyses and PCoA plots provided further evidence that ATMs altered gut microbiome structure, and also demonstrated that ATMs introduced significant variation within microbial communities. These effects did not differ across sex, except microbiomes of ATM females showed greater reductions in rare species. Unexpectedly, pre-ATM-treated mice showed a difference in weighted UniFrac distances relative to controls. Given the absence of differences in all other measures on Day 0, our small sample size at this timepoint, and the randomization of previously group-housed subjects into ATM and control groups, it is unlikely that this difference is meaningful. Our findings that no measures of alpha or beta diversity differed across treatment Days 3, 7, and 10 suggest that our ATM cocktail alters gut microbiome structure in a uniform manner for at least one week. Given that microbiomes of controls did not differ across these days either, these findings also suggest that the behavioral tests our mice were exposed to were not high-stress experiences that induced gut dysbiosis on their own.

Taxonomic differences between ATM mice and controls were apparent by Day 3 of treatment and did not differ across time in either sex, providing further support that ATMs induce a relatively stable state of gut dysbiosis for at least one week. The largest shifts included a reduction in Firmicutes, especially Clostridiales, and a relative increase in Verrucomicrobia and Proteobacteria, especially Gammaproteobacteria. We also identified a large sub-community of Clostridia and Bacteriodetes species whose dominance was reduced by ATM treatment. Identification of the species most relevant to the effects observed in this study will require further investigation. We note that, while our approach does not encompass other members of the gut microbiome (viruses and eukaryotes), bacteria are the most likely source of neuroactive components and metabolites.

Taken together, our data demonstrate that our ATM cocktail induced gut dysbiosis in both females and males, and that these dysbiotic states were at least partly distinct. The larger impact of ATMs on phylogenetic diversity, sub-communities, and rare species in females relative to males suggests that there were sex-specific gut microbiome responses to treatment. Whether the sex-specific effects ATMs exerted on animal behavior and BLA dendritic spine density in this study were a direct consequence of changes in gut microbiota or a more downstream effect due to other biological differences in the way females and males respond to gut dysbiosis remains unclear.

### 4.6 Limitations of ATM Treatment

While ATM models of gut dysbiosis are plentiful, treatment regimens (ATMs used, doses, and lengths of administration) vary widely, and strain-dependent dosing remains to be established (Bercik et al., 2011; Jang, Lee, Jang, Han, & Kim, 2018; Kennedy, King, & Baldridge, 2018; Odeh, 2013). To our knowledge, three other studies have used our cocktail to study the MGB axis. Based on findings that higher doses may deleteriously affect consumption (Odeh, 2013), we used a dose that was relatively low compared to the other two studies (Bercik et al., 2011; Verdu et al., 2006). Nonetheless, treatment reduced body weight and water consumption in both sexes across multiple days (Figure S1, S2). Some studies circumvent this issue through the use of nutritive and non-nutritive sweeteners, but these additives limit interpretability of results by altering reward circuitry and gut microbiome composition (Ruiz-Ojeda et al., 2019). Although we observed no declines in animal health, weight loss remained under 15%, and water consumption mildly reduced only on some days in ATM females, we questioned whether decreases in drinking behavior could have affected our results. Although this possibility cannot be ruled out, it does not seem likely. Water deprivation does not affect aversive learning or cued recall in studies with comparable methodologies (3 CS/US pairings; Maren, DeCola, & Fanselow, 1994; Pouzet, Zhang, Richmond, Rawlins, & Feldon, 2001). Although water deprivation is sometimes associated with hyperlocomotor activity, this phenotype is inconsistently reported, and tends to emerge only in the dark cycle (Tsunematsu et al., 2008). Furthermore, if deficits in cued freezing were caused by changes water consumption or body weight, these results should have been apparent in the second experiment as well as the first.

### 4.7 Mechanistic underpinnings of ATM-induced behavioral phenotypes

MGB signaling occurs through multiple pathways, primarily neural, immune, and endocrine, which are bidirectional and interactive (El Aidy, Stilling, Dinan, & Cryan, 2016; Forsythe, Kunze, & Bienenstock, 2016; Martin, Osadchiy, Kalani, & Mayer, 2018). A higher dose of our ATM cocktail was found to decrease anxiety-like behavior and amygdala BDNF in male mice independent of vagal and sympathetic integrity, and in the absence of changes in inflammatory profile, intestinal morphology, and specific enteric neurotransmitters; this suggests that other microbial metabolites were responsible for the observed effects (Bercik et al., 2011). Our cocktail induced distinct shifts in gut microbiome composition relative to those in Bercik et al. (2011), but at least partially shared underlying mechanisms are likely. Another notable difference is that we studied both females and males, and observed effects of sex and, preliminarily, estrous stage. The relative impact gonadal hormones had on our data is unclear, but it is notable that gonadal hormones share numerous reciprocal connections with gut microbiota (Baker, Al-Nakkash, & Herbst-Kralovetz, 2017; Clarke et al., 2013; Tetel, De Vries, Melcangi, Panzica, & O’Mahony, 2018) and modulate amygdala activity in both rodents and humans (Blume et al., 2017; Velasco, Florido, Milad, & Andero, 2019; Womble, Andrew, & Crook, 2002). Overall, the parallel and sex-specific pathways by which our cocktail impacts the central nervous system require further study.

### 4.8 Gut Microbiome and the Basolateral Amygdala

Our findings have important implications at the level of the BLA. One explanation for the female-specific decreases in conditioning performance and BLA dendritic spine density in this study is that ATMs differentially altered female and male BDNF-TrkB signaling. It is well-established that BDNF is sensitive to perturbations of the gut microbiome (Clarke et al., 2013; Heijtz et al., 2011; Hoban et al., 2016; 2017; Sudo et al., 2004), and our cocktail has been linked to decreased amygdala BDNF (Bercik et al., 2011). Given that binding of BDNF to the TrkB receptor in the BLA is necessary for the acquisition and consolidation of aversive memory (Rattiner, Davis, French, & Ressler, 2004; Rattiner, Davis, & Ressler, 2005), it is possible that females are particularly susceptible to effects of gut dysbiosis on BLA BDNF. Indeed, accumulating data indicates that there are sex differences in stimulus-induced *BDNF* expression in various brain regions, including the amygdala, as well as sexspecific variations in TrkB signaling (see Chan & Ye, 2017 for a review).

Although it is unlikely that peripheral inflammatory processes played a role in our findings, sex differences in microglial activation are another noteworthy potential mechanism. Microglia are highly heterogenous in structure and function: they are sexually differentiated, responsive to reproductive hormones, and actively regulate nearly all aspects of neurotransmission (Lenz & Nelson, 2018; Tay, Savage, Hui, Bisht, & Tremblay, 2017). Microglia also express TrkB and release BDNF themselves, and microglia-depleted mice show reduced freezing in auditory cued aversive recall (Parkhurst et al., 2013; Spencer-Segal et al., 2011). Additionally, impaired aversive extinction in adult ATM-treated and GF male mice is linked with altered microglial expression (Chu et al., 2019). Much about the interactions between sex, BNDF-TrkB signaling, and microglia remain elusive, but these are promising areas for exploration.

### 4.9 Clinical Implications

Trauma and anxiety-related disorders are major health burdens worldwide, with increased incidence in women relative to men (Altemus et al., 2014). Associated symptoms are highly prevalent in patients with dysbiosis-related syndromes like IBS, which also disproportionately affect women (Kibune-Nagasako et al., 2016; Lovell & Ford, 2012). Our data add to growing evidence for a role of gut microbiota in the pathophysiology of psychiatric conditions (Cowan et al., 2018; Cryan & Dinan, 2012; Kelly et al., 2016; Malan-Muller et al., 2018; MacQueen, Surette, & Moayyedi, 2017), and highlight the need to consider sex as a biological variable (Shansky & Woolley, 2016). While animal models can provide powerful insight into mechanisms underlying disease states, caution should be exercised in translating results. MGB research is still in its infancy, and “dysbiotic patterns” remain poorly defined, and differ between species as well as conspecifics (Walter, Armet, Finlay, & Shanahan, 2020). Nonetheless, elucidation of the effects observed in this study could have important clinical utility moving into the future.

### 4.10 Conclusions

Our data bridge a research gap in the characterization of sex-dependent influences of gut microbiota on various nodes of aversive conditioning and anxiety-related circuitry. Specifically, we found novel evidence that female mice are more susceptible than males to microbial modulation of aversive learning. This was supported by sex-specific effects of ATM treatment on BLA dendritic spine patterning. We also found that, while females are more susceptible than males to ATM-induced alterations in gut microbial communities, their anxiety-like behaviors are more robustly buffered. These findings emphasize the necessity of studying both sexes and advance our understanding of the broad spectrum of influences gut dysbiosis can exert during adulthood.

## AUTHOR CONTRIBUTIONS

All authors read and approved the final manuscript. *Conceptualization*, C.G.G.; *Methodology*, C.G.G., H.C.B., D.J.E; V.C.W. *Investigation*, V.C.W., K.L.B., P.O.J., T.B., V.B., C.G.G., C.B.B., A.F.P., A.M., H.M.T., and S.R.; *Formal Analysis*, H.C.B., D.J.E., J.C.T, V.C.W., and C.G.G.; *Resources*, H.C.B. and D.J.E; *Writing-Original Draft* and *Review and Editing*, C.G.G., H.C.B., D.J.E., J.C.T., and V.C.W.; *Visualization*, H.C.B., D.J.E., V.C.W, and C.G.G.; *Supervision*, H.C.B. and D.J.E.; *Funding Acquisition*, H.C.B., D.J.E., and C.G.G.

## OTHER CONTRIBUTIONS

The authors are grateful to Vassar College Professors Kelli Duncan and Bojana Zupan for their guidance on estrous stage determination and brain tissue processing, respectively. This study was supported by grants from the Asprey Center for Collaborative Approaches in Science, the Lucy Maynard Salmon Research Fund at Vassar College, and the Ruth M. Berger Foundation.

## DATA ACCESSIBILITY LINKS

The microbiome sequence data in this study will be made available at the Sequence Read Archive (SRA) under BioProject ID: PRJNA646617. The BLA principal neuron dendrite reconstructions will be made available at NeuroMorpho.org. Other data that support our findings are available from the corresponding author upon reasonable request.

