## Supplemental Figures 1, 2, 3, and 4 for "Sex-specific gut microbiota modulation of aversive conditioning and basolateral amygdala dendritic spine density"

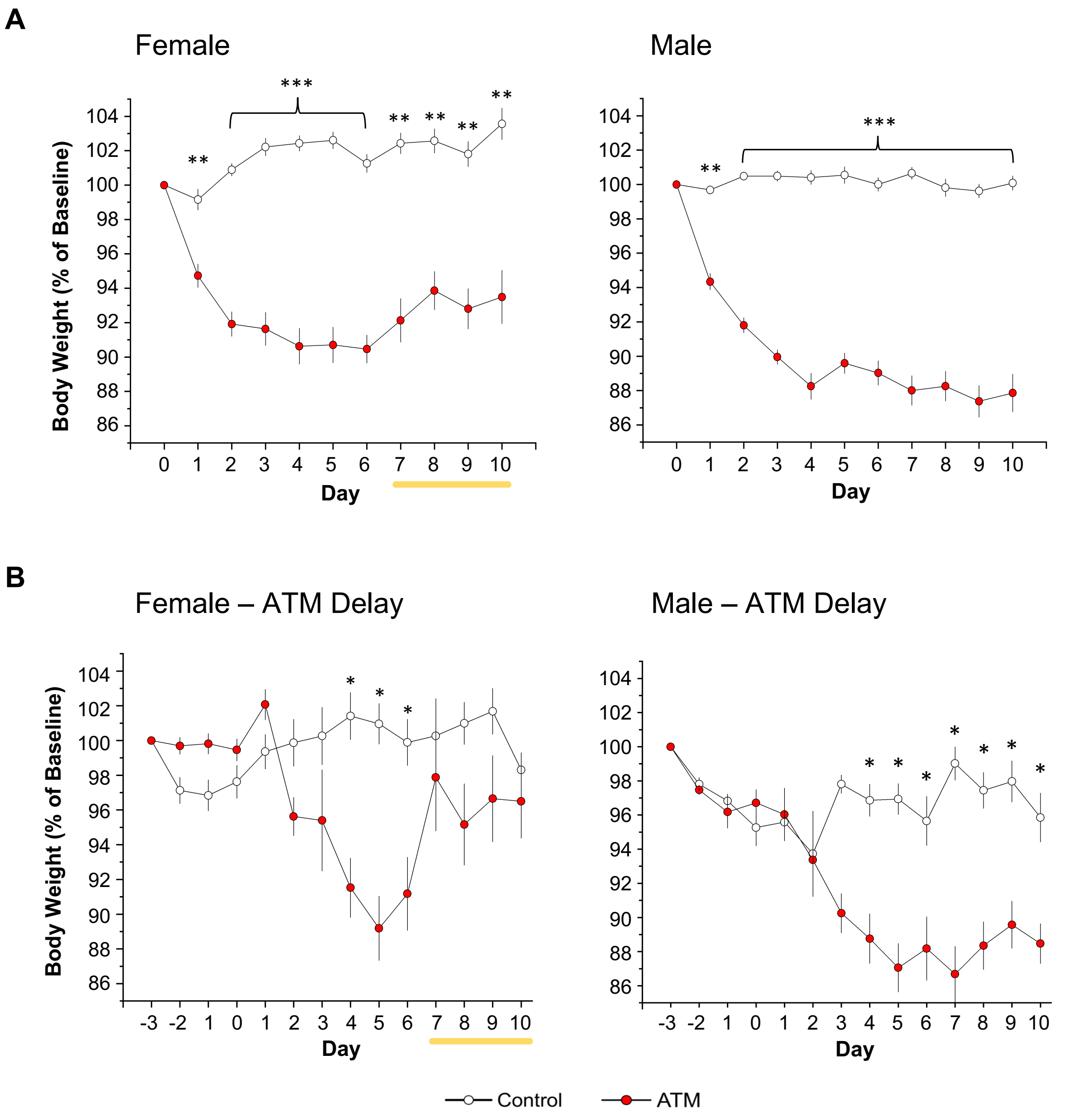


**Figure S1. ATMs decrease body weight in both sexes.** (***A)*** Relative to controls, treatment reduced body weight in both females (n = 23/group) and males (n = 24/group) across Days 1–10. Only females showed a rebound in body weight across Days 7–10. ***(B)*** In the ATM Delay experiment, treatment reduced body weight in both females (n = 12/group) and males (n = 12/group) starting on Day 4. Consistent with the first experiment (and emphasized by the yellow lines), females showed a rebound in body weight across Days 7–10. **P* < 0.05, ***P* < 0.01, ****P* < 0.001

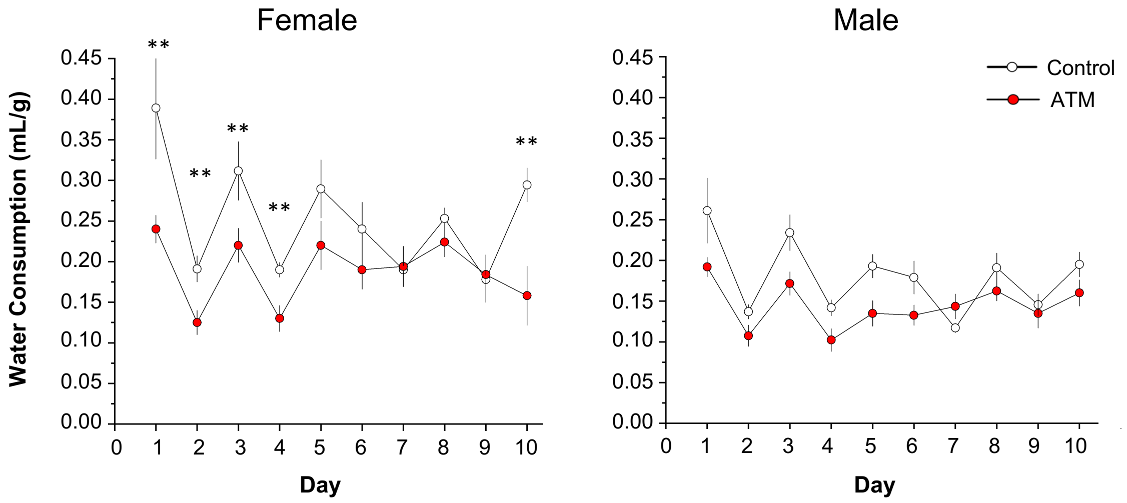


**Figure S2. ATMs mildly attenuate voluntary water consumption in females but not males.** Because ATMs were administered in drinking water, we assessed each mouse's daily water bottle volume loss to determine whether treatment influenced voluntary water consumption. Since average fluid intake can differ across body size, we normalized data by dividing each subject's water bottle loss (mL) by its daily weight (g). ATM females lost more drinking water than female controls on Days 1–4, and on Day 10 (n = 23/group), suggesting that treatment mildly reduced drinking behavior. ATMs did not affect water consumption in males (n = 24/group). *$P$ < 0.05

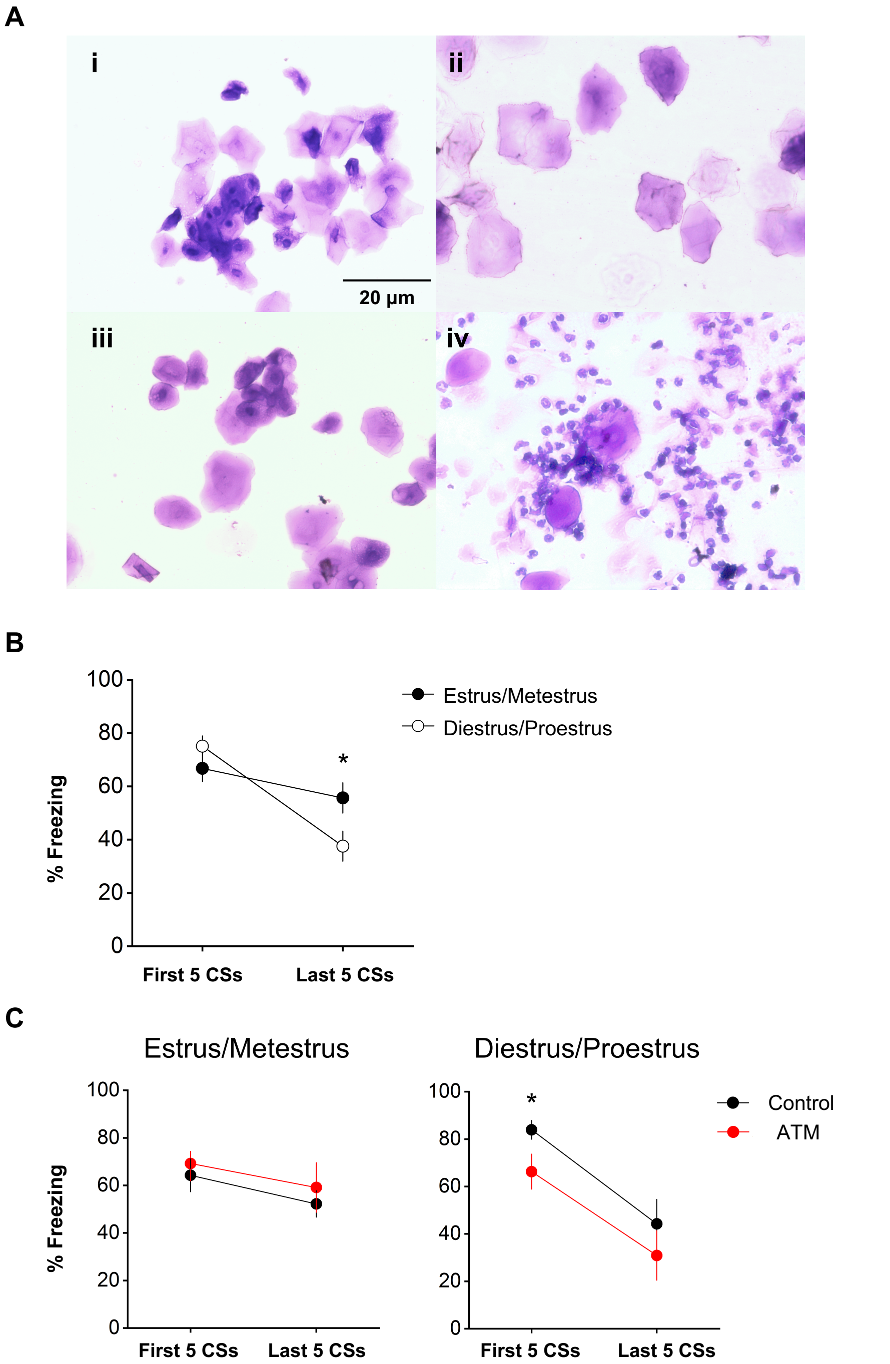


**Figure S3. Ovarian hormones modulate aversive extinction and interact with ATMs to influence cued recall**. ***(A)*** Micrographs (40X) of 0.1% crystal violet-stained vaginal cells during *(i)* proestrus, *(ii)* estrus, *(iii)* metestrus, and *(iv)* diestrus. ***(B)*** Mice in diestrus/proestrus showed less freezing than mice in estrus/metestrus during the final 5 presentations of the CS, suggesting that high levels of ovarian hormones attenuate aversive extinction. There was no main effect of estrous stage on cued recall. ***(C)*** ATM mice in diestrus/proestrus, but not estrus/metestrus, showed less freezing relative to controls during the first 5 presentations of the CS, suggesting that gut dysbiosis interacts with high levels of ovarian hormones to attenuate retrieval of an aversive CS. n = 6/group, $^*P < 0.05$

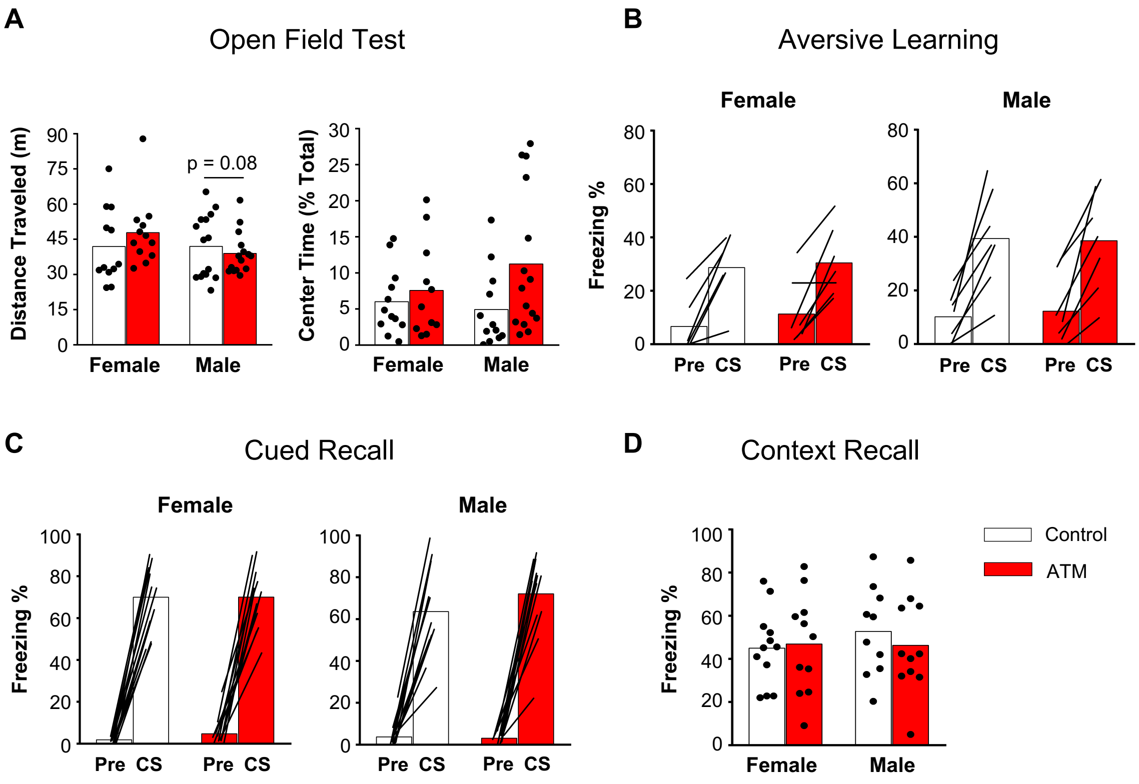


**Figure S4. ATMs do not alter cued expression in either sex during the ATM Delay experiment.** ***(A)*** Locomotor activity did not differ between females and males and was not affected by ATMs in either sex. Percentage time spent in the arena's center trended towards an increase in ATM males but did not differ between female groups (n = 12/group). ***(B)*** Groups did not differ in aversive learning (n = 6–7group), ***(C)*** cued recall (n = 12/group), or ***(D)*** contextual recall (n = 10–12/group). *$P < 0.05$
